# Deciphering the gene regulatory circuitry governing chemoresistance in Triple-Negative Breast Cancer

**DOI:** 10.1101/2023.05.05.539623

**Authors:** Ryan Lusby, Ziyi Zhang, Arun Mahesh, Vijay K. Tiwari

## Abstract

Triple-negative breast cancer (TNBC) is the most aggressive breast cancer subtype, due in part to extensive intratumoral heterogeneity, high rates of metastasis and chemoresistance, leading to poor clinical outcomes. Despite progress, the mechanistic basis of chemotherapy resistance in TNBC patients remains poorly understood. Here, using single-cell transcriptome datasets of matched longitudinal TNBC chemoresponsive and chemoresistant patient cohorts, we discover cell subpopulations associated with chemoresistance and the signature genes defining these populations. Notably, we show that the expression of these chemoresistance genes is driven via a set of TNBC super-enhancers and transcription factor networks across TNBC subtypes. Furthermore, genetic screens reveal that a subset of these transcription factors is essential for the survival of TNBC cells and their loss increases sensitivity to chemotherapeutic agents. Overall, our study has revealed transcriptional regulatory networks underlying chemoresistance and suggests novel avenues to stratify and improve the treatment of patients with a high risk of developing resistance.

## INTRODUCTION

Triple-negative breast cancer (TNBC) is a highly heterogeneous disease defined by the absence of oestrogen receptor (ER) and progesterone receptor (PR) expression and human epidermal growth factor receptor 2 (HER2) overexpression ^1^. It is associated with a poorer clinical outcome due to a lack of early prognostic techniques, high incidences of relapse, metastasis and a lack of targeted therapeutics ^2^. In the neoadjuvant setting, chemotherapy is the standard treatment, which includes a combination of taxanes and anthracyclines. However, approximately 30%-50% of patients develop resistance, and their prognosis worsens to 13-15 months survival ^3, 4^. Despite TNBC being grouped as a single disease, clinical, histological, and molecular profiling have highlighted its intrinsic heterogeneity ^5^. This heterogeneity is further highlighted with the identification of unique TNBC subtypes (TNBC type-4 classification) that include: basal-like 1 (BL1), basal-like 2 (BL2), mesenchymal (M) and luminal androgen receptor (LAR) ^6^. Each subtype displays unique transcriptional patterns, biology and chemotherapy response ^7, 8^.

The distal gene regulatory landscape plays a critical role in driving disease-associated altered cell-fates^9^. A super-enhancer (SE) is a cluster of enhancers initially found to be essential in determining cell identity during differentiation but have progressively been implicated in disease initiation and progression, including tumorigenesis ^10–12^. In breast cancer, it has been demonstrated that enhancer and SE transcription can reveal insights into subtype-specific gene expression programs ^13^. SEs exhibit high transcription factor density, especially for Core Regulatory Circuitry (CRC) transcription factors (TFs) and drive the expression of key genes that strongly influence cellular identity and function ^11, 14, 15^. These CRC TFs have been shown to self-regulate, where they inwardly bind to SE regions and outwardly regulate SE-associated genes with the CRC, forming a forward-feeding loop. Accordingly, disrupting SE structure or inhibiting SE targeting factors has shown promising results as a potential therapeutic avenue for certain cancers ^16, 17^. Surprisingly, however, the contribution of SEs and associated CRC landscapes in regulating the gene regulatory programs underlying TNBC aggressiveness remains unknown. In particular, it remains to be known whether TNBC subtype-specific super-enhancers and CRCs exist to confer different degrees of chemoresistance in these subtypes.

In this study, we aimed to address these longstanding questions by characterising the epigenomic, transcriptomic and TF landscape underlying chemoresistance in TNBC patients. By profiling matched longitudinal single-cell RNA-sequencing data (scRNA-seq) of chemoresponsive and chemoresistant TNBC patients, we identified unique cellular subpopulations associated with chemoresistance and revealed genes that define these subpopulations. Notably, a subset of these signature genes outperformed existing gene panels in classifying pathologic complete response versus persistent residual disease against pre-operative neoadjuvant chemotherapy in TNBC. Furthermore, by analysing data from H3K27ac Chromatin immunoprecipitation followed by sequencing (ChIP-seq) of TNBC subtype patients, we define the SE architecture and CRCs associated with the gene expression programme underlying chemoresistance and reveal several TFs whose depletion can improve the efficacy of chemotherapy across TNBC subtypes.

## RESULTS

### A subpopulation specific gene expression signature associates with aggressiveness in chemoresistant TNBC patients

We began by outlining a stepwise plan to reveal the gene regulatory circuitry underlying chemoresistance in TNBC patients (Fig. 1A). Due to the high degree of intra-tumour heterogeneity associated with TNBC, scRNA-seq provides a higher level of resolution and enables the identification of minor changes in gene expression profiles within tumour cells, being embedded with multiple cell types in a varying proportion which could be lost in bulk RNA-seq analysis. To unravel the underlying mechanisms of chemoresistance in TNBC, we obtained and analysed a scRNA-seq dataset consisting of a total of 6,862 cells from four responsive (chemosensitive) and four resistant (chemoresistant) patients to neoadjuvant chemotherapy (NAC) at pre-and post-treatment time points, containing only tumour cells as they were pre-selected before sequencing (Supplementary Table 1). To classify the tumours as sensitive or resistant in the scRNA analysis, Kim et al^18^ had performed deep-exome sequencing on 20 patients in which they identified 10 patients where NAC led to clonal extinction (sensitive) and 10 patients where clones persisted (resistant) after treatment. From these 20, 8 patients (4 sensitive and 4 resistant) were selected for single-cell RNA sequencing^18^. We hypothesized that by profiling TNBC chemoresistant patient data at the single-cell level, we could identify critical markers driving chemotherapy response. Furthermore, identifying these markers could enable the prediction of chemotherapy response in untreated patients.

**Figure 1.**
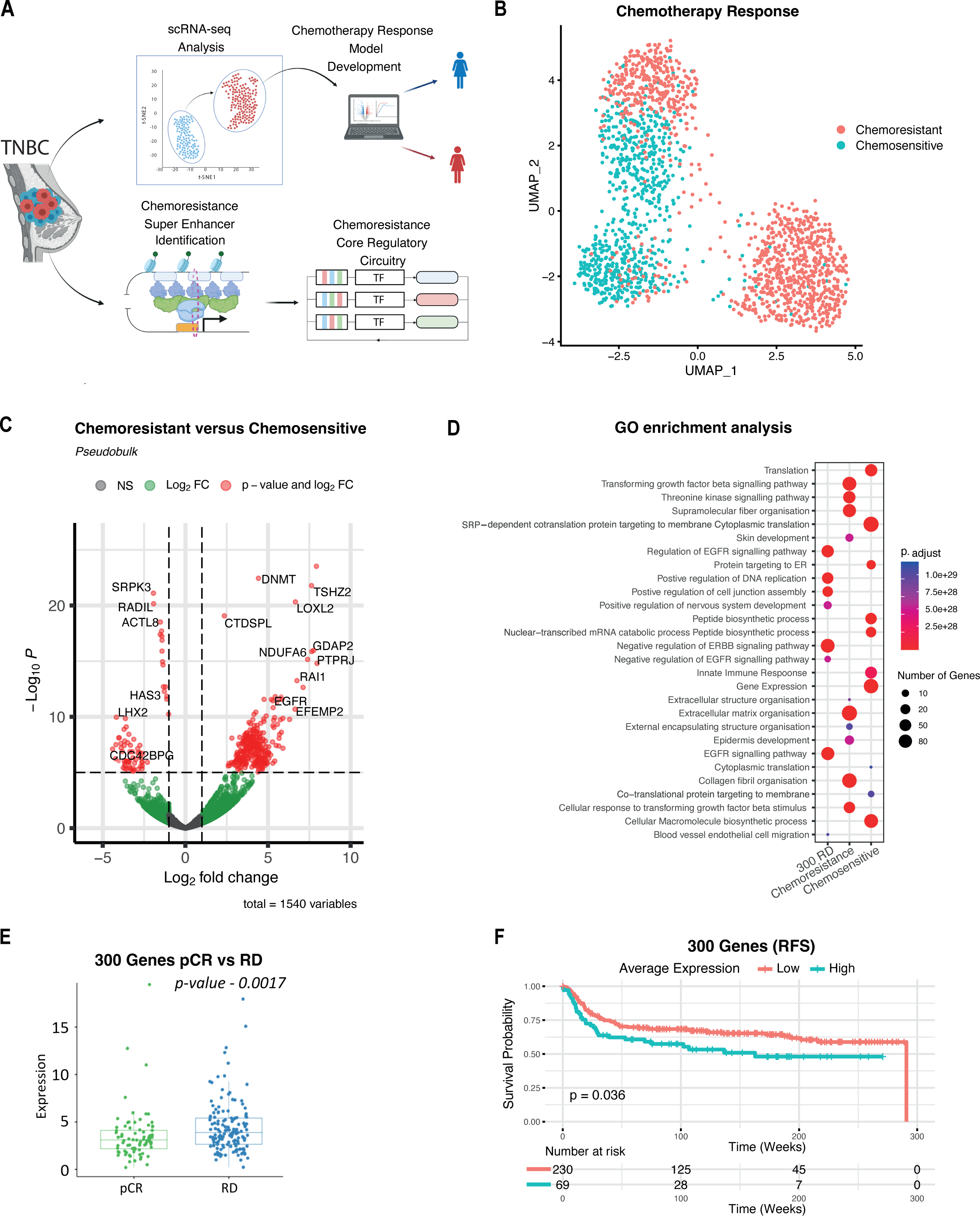
scRNA-seq Analysis Reveals Subpopulations of Cells and Key Genes Underlying TNBC Chemoresistance. **A)** Graphical abstract of study workflow **B)** UMAP projection of pre-treatment chemoresponsive and chemoresistant patients **C)** Volcano plot of differentially expressed genes highlighting key transcriptional differences between chemosensitive and chemoresistant patients. **D)** Gene Ontology analysis of markers for pre-treatment chemoresistant and chemoresponsive patients and the 300 genes reveals they are significantly involved in signalling pathways and cell migration. **E)** Reproducibility analysis reveals that 300 markers identified in pre-treatment chemoresistant patients had higher expression in residual disease than pathologic complete response across all datasets (Wilcoxon rank-sum test p=0.0017). **F)** Survival plot of the 300 genes in TNBC patients from the METABRIC Cohort (Kaplan–Meier, p=0.036).

As it is thought that chemoresistance occurs due to the clonal evolution of pre-existing clones ^18, 19^, we focused our analysis on markers unique to the pre-treatment chemoresistant patients to identify the critical transcriptional landscape defining chemotherapy response. In the original study^20^, the authors had highlighted that the batch effects were minimal between patient samples and hence we were convinced we could proceed with merging. By merging scRNA-seq data from all 8 patients, we extracted the cells of the pre-treatment samples (Fig. 1B, Supplementary Fig. 1A-D). Clustering analysis of pre-treatment cells revealed that chemoresistant and chemosensitive patients had overlapping clusters, highlighting the lack of batch effects identified by Kim et al, but also a distinct, separate cluster of chemoresistant cells, highlighting a subset of cells that may play a role in patients showing a poor response to chemotherapy (Fig. 1B). Cell annotation analysis, performed by SCSA using established cell type markers from two public databases: CellMarker and CancerSEA. database^21^, revealed that chemoresistant clusters were predominately basal epithelial cells whilst chemosensitive clusters contained luminal progenitor and luminal epithelial cells (Supplementary Fig. 1B). Interestingly, progenitor cells are more likely to be sensitive to anti-cancer therapies^22^, whilst luminal epithelial cells can give rise to basal epithelial cells upon oncogenic stress^23^. To gain an insight into the transcriptional landscape driving chemoresistance, in pre-treatment patient samples, we applied pseudobulk differential gene expression analysis between chemosensitive and chemoresistance annoataions. We identified distinct and statistically significant gene expression patterns for each condition (p-value >=0.05 and logfc>=1) (Fig. 1C, Supplementary Fig.1E). Gene ontology analysis showed enrichment of extracellular matrix remodelling and transforming growth factor-beta (TGF-β) signalling (Fig. 1D), processes associated with EMT, confirming the results from Kim et al, and which have previously been implicated in TNBC chemoresistance^24^. Together these results highlight the existing differential transcriptional landscape of chemoresistant and chemosensitive TNBC patients prior to NAC treatment.

Due to the low patient numbers in the scRNA-seq data, we next sought to identify genes with a reproducible expression in a larger cohort of patients. To address this issue, we obtained and processed bulk RNA-seq datasets (GSE20271, GSE25055, GSE25065, GSE20194 and GSE163882) consisting of 397 TNBC patients, pre-NAC, with known outcomes of pathologic complete response (pCR) and residual disease (RD). To ensure that batch effects between studies were minimal, we corrected using the established R package SVA and the function ComBat which uses empirical Bayes frameworks for adjusting data for batch effects^25^. We found that there were very little batch effects following merging that were corrected post-analysis (Supplementary Fig. 2A-B). To assess the reproducibility of the genes in a larger patient cohort we compared expression levels of each gene between RD and pCR across 397 TNBC patients total. This resulted in the identification of 300 marker genes which showed a significantly higher expression across all patients with RD (Fig. 1E). By implementing Kaplan– Meier estimator survival analysis on RNA-seq data from the TNBC METABRIC cohort, we revealed that high average expression of these 300 genes is associated with a significantly decreased relapse-free survival in TNBC patients whilst using the median expression as the cut-off point to stratify patients into high and low subgroups (Fig 1F). In addition, following the reduction in gene numbers due to many having non-detectible expression in bulk RNA-seq data, Gene Ontology analysis revealed that these genes were significantly involved in EGFR signalling pathway (Fig 1D.), which is previously implicated in TNBC chemoresistance ^26–28^.

### Distinct transcription factor regulons are active in pre-treatment chemoresistant cells

Currently the regulatory landscape driving TNBC chemoresistance is unknown. We sought to address this by investigating potential regulatory mechanisms which govern expression of chemotherapy resistant genes, we deployed a single-cell regulatory network inference and clustering (SCENIC) ^29^ computational pipeline to identify regulons (TFs and their targets) and assess their activity in the chemoresistant cell populations compared to chemosensitive ones (Fig. 2A). In brief, first co-expression modules were identified using GRNBoost. Next, the motifs driving resistant cells were discovered using cisTarget. Finally, the regulon activity was quantified by assessing enrichment of the regulon target genes using AUCell^30^. Through this analysis, we identified regulons with high activity and specificity scores for both chemoresistant and chemosensitive cells (Fig. 2B-C, Supplementary Table 2). Of note, the Transcription Factor (TF) TFAP2C was identified among the top regulons based on the AUCell score in chemoresistant cells and not present in chemosensitive cells (Fig. 2C-D) and has previously been implicated in EMT signalling and chemoresistance in lung adenocarcinoma, but not yet in TNBC ^31, 32^. Additionally, we discovered SP1 which was shown to promote chemoresistance and metastasis in ovarian cancer and breast cancer^33^. In both cases, it has been implicated in EGFR transactivation and facilitating migration and invasion through Smad3 and ERK/Sp1 signalling pathways ^33, 34^. Furthermore, another regulon TFAP2A has also have been associated with chemoresistance in colorectal cancer but not yet in TNBC ^35^. Interestingly, expression of many of these TFs including TFAP2C, TFAP2A and SP1 were higher in treatment naïve chemoresistance patients as compared to the chemoresponsive patients and these patterns persisted post-chemotherapy (Fig. 2E). Such higher expression and activity of these TFs in resistant patients compared to sensitive implicates these TFs among the key contributors of chemoresistance in TNBC patients.

**Figure 2.**
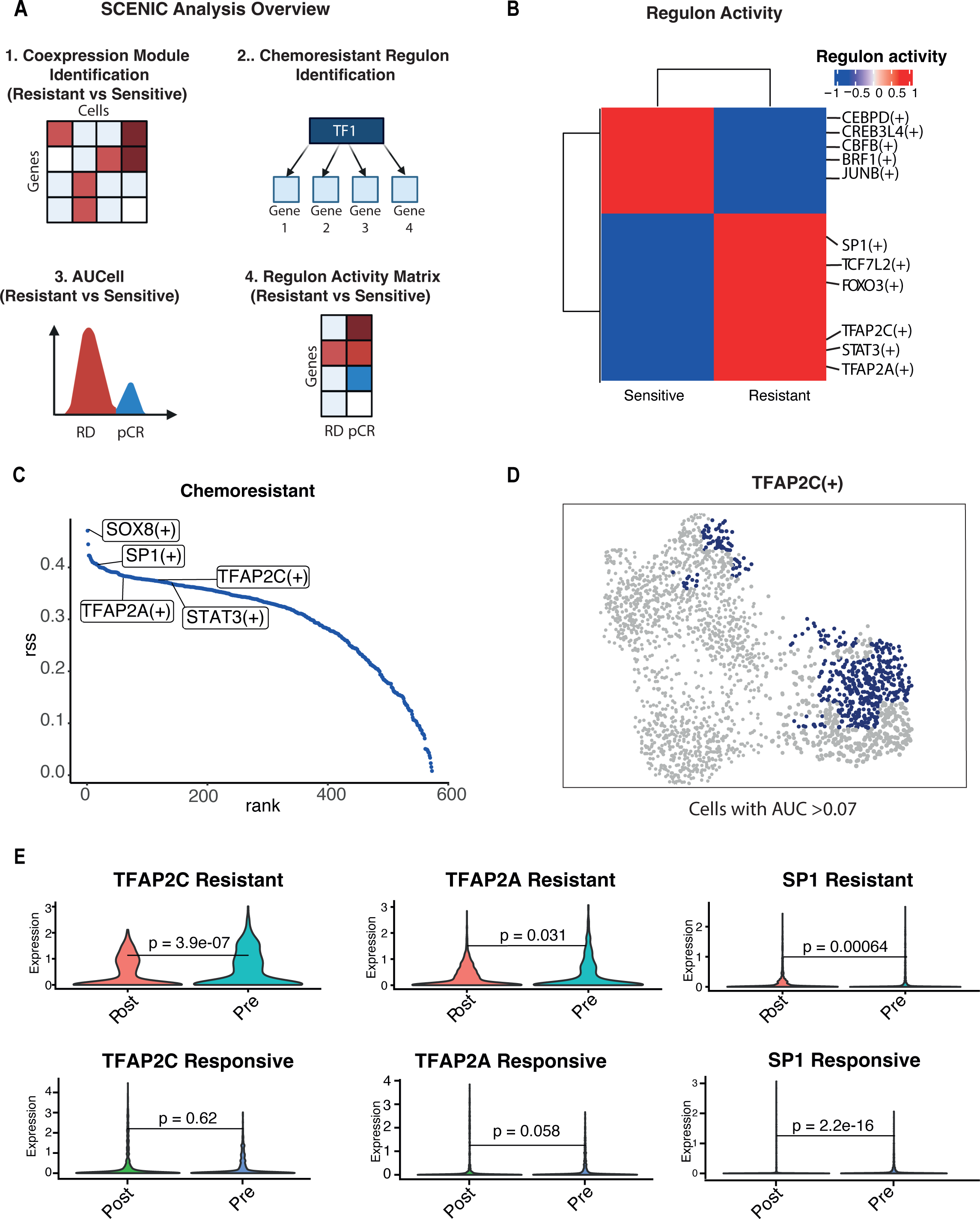
SCENIC analysis reveals potential chemoresistance gene regulons. **A)** SCENIC Workflow: 1) Identification of co-expression modules between resistant and sensitive cells using GRNBoost 2) Regulon identification using cisTarget 3-4) the regulon activity was quantified by assessing the enrichment of the regulon target genes using AUCell. **B)** Heatmaps show significant top regulators based on AUC score for chemosensitive and chemoresistant patients, top motifs are labelled (Binary Score). **C)** RSS Plot showing top regulons for chemoresistant clusters with top motifs highlighted. **D)** UMAP highlighting cells with AUC >0.07 TFAP2C regulon activity. **E)** Violin plots of TFAP2C, TFAP2A and SP1 expression in Resistant and Responsive patients scRNA-seq highlighting that higher expression remains following treatment in chemoresistant patients (Wilcoxon rank-sum test).

### A minimalistic gene signature of 20 genes can predict chemotherapy response in treatment naïve TNBC patients with high accuracy

In TNBC, patients achieving a pathologic complete response to neoadjuvant chemotherapy is a crucial predictor of a patient’s long-term outcomes and can allow an early evaluation of the effectiveness of systemic therapy^3, 36^. We next wanted to investigate whether the genes identified are potentially the critical drivers of chemoresistance in these patients by identifying a significant gene set that could accurately stratify RD and pCR patients. By utilising Lasso and Elastic-Net Regularized Generalized Linear Models, we identified a significant gene set that could accurately differentiate between pCR and RD in TNBC patients. We derived our training dataset by combining GSE20271 and GSE25055 datasets with 177 TNBC patients (57 pathologic complete response, 120 residual disease), and to derive the validation dataset, we combined GSE25065 and GSE20194 datasets with 130 TNBC patients (46 pathologic complete response, 84 residual disease). We combined these datasets to increase the training and testing cohorts to improve the strength and validity of the proposed gene model as previously shown^37^. In brief, we built a single-fold lasso-penalised model for all genes, in the training dataset, then performed 10-fold cross-validation (Supplementary Fig. 3A-B) to identify the best predictors of RD vs pCR. We then took our top predictors (Fig. 3A), built a new model and performed ROC analysis on our validation dataset. This analysis revealed a total of 20 genes (CLCN3, NDUFA6, PTPRJ, GDAP2, RNF19B, MKKS, TSHZ2, COL21A1, LOXL2, SLC11A2, ESM1, CTDSPL, RAI1, EFEMP2, DTNA, EPHB3, EGFR, HOXA1, MSH3 and PPFIA2) to have the strongest discriminatory power between RD and pCR, training AUC = 0.90 (Fig. 3B), Validation AUC = 0.89 (Supplementary Fig. 3C).

**Figure 3.**
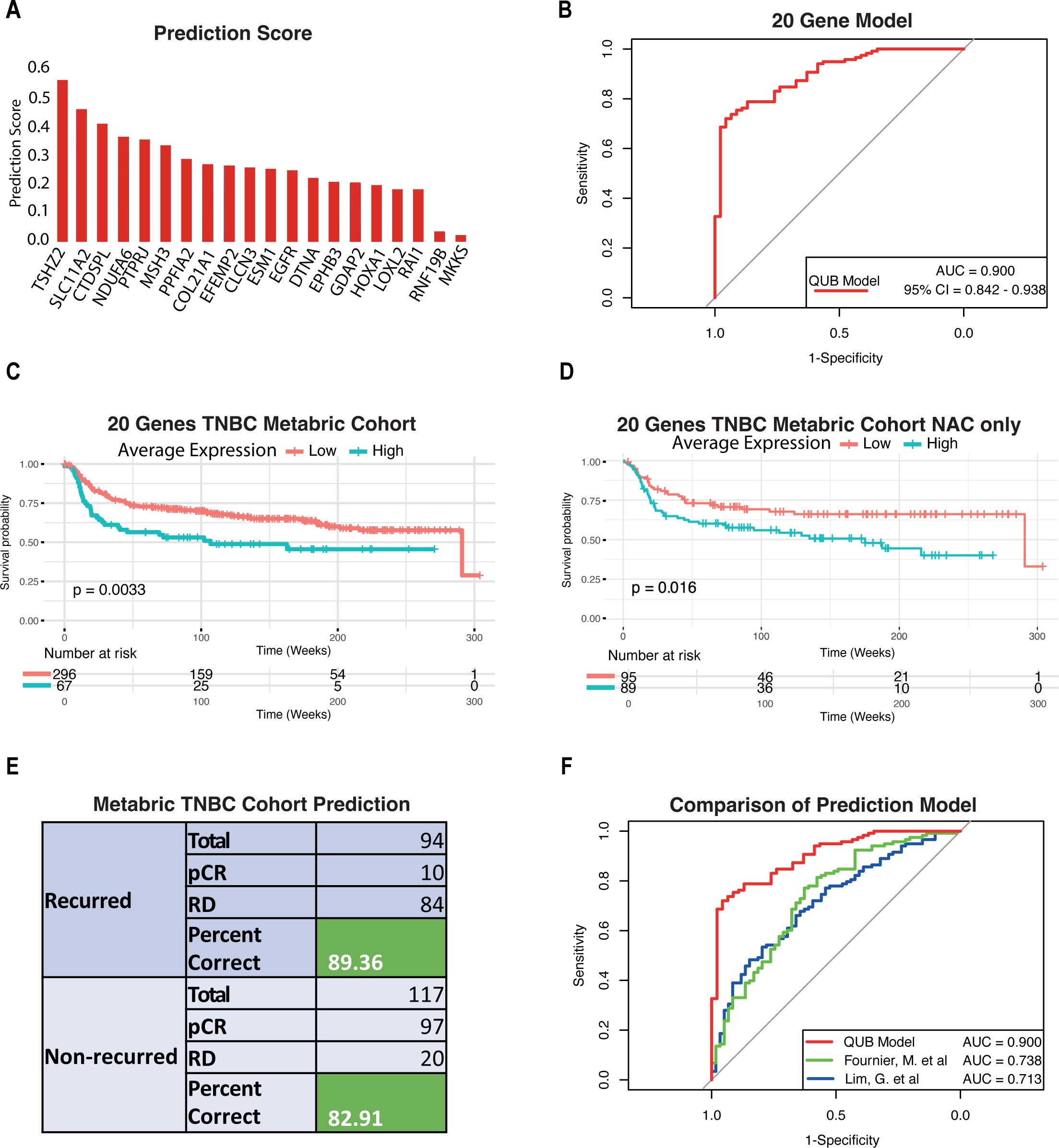
20 Gene Models shown to have a high accuracy in predicting chemotherapy response in TNBC patients. **A)** Ranked score of each gene in determining RD **B)** ROC curve highlighting the accuracy of our model **C)** Survival plot of the 20 genes in TNBC patients from the METABRIC Cohort (Kaplan–Meier, p=0.0033) **D)** ROC curve highlighting the accuracy of our model compared to previously published TNBC prediction models. **E)** Survival plot of the 20 genes in TNBC patients from the METABRIC Cohort who received NAC only (Kaplan–Meier, p=0.016) **F)** Prediction of each TNBC METABRIC patients based on relapse free survival.

As we have been focusing on TNBC only, we next sought to explore the role of the 20 gene model in other breast cancer subtypes. We first investigated the the expression of all 20 genes in the TCGA-BRCA dataset containing the four primary subtypes of breast cancers. In combination, the average expression of the 20 genes showed significantly higher expression in the Basel subtype compared to Luminal A and B breast cancer subtypes but lower in the HER2 subtype (Supplementary Fig. 3D). We next wanted to expand the utility of our 20 gene panel to include a prognostic capability through testing 5 year relapse free survival. Notably, survival analysis of all TNBC patients from the METABRIC cohort, using median expression as the cut-off value, revealed that in combination high expression of this gene signature is associated with significantly reduced relapse-free survival over five years (Fig. 3C). However, in luminal A/B and HER2 patients’, higher expression of the gene set had no correlation with increased or reduced survival (Supplementary Fig. 3E-F). Highlighting that higher expression of these genes in TNBC only, is underlying their chemoresistance potential. Furthermore, when filtering TNBC METABRIC patients for those only receiving NAC we found that high expression of the 20 genes is again associated with reduced survival irrespective of the chemotherapy regimen (Fig. 3E). Altogether, these findings suggest that increased expression of these genes in treatment naïve TNBC patients may drive chemoresistance leading to poor outcomes.

Whilst we had built and tested the model on two large cohorts we next sought to further validate our model’s predictive strength by applying it to the TNBC METABRIC cohort. Again, whilst not considering which chemotherapy regime was applied and using patients’ relapse-free status as a determination of pCR and RD, our model successfully predicted 89.4% of RD and 82.9% of pCR patients correctly (Fig. 3E). This outcome successfully highlights, not only the predictive strength of our model but also highlights that high expression of the 20 genes can give insights into patients relapse free survival. Unlike other breast cancer subtypes, there are currently no tests in clinical use for TNBC patients to accurately predict NAC response and facilitate their clinical management ^38^. While several predictive panels have been published for TNBC, none have achieved clinical utility due to small sample sizes, lack of validation data and inability to achieve the necessary predictive strength in oestrogen and HER2 positive tumours ^7, 37, 39–41^. Addressing this critical unmet need, we show that our 20 genes hold a high predictive power in determining RD or pCR (with an area under the curve (AUC) of 0.90) (Fig. 3B). Notably, our model utilising this minimal 20 gene set outperformed all previous models in predicting chemotherapy response in TNBC^37, 41^ (Fig. 3F). The specific combination of these 20 genes was essential for its high performance as further removal of genes (top 5) significantly reduced the accuracy (AUC: 0.827) (Supplementary Fig. 3G). Additionally, by applying 20 cross-validation iterations we found that the predictive strength remained the same (Supplementary Fig. 3H).

One of the key factors that result in TNBC being the most aggressive breast cancer subtype is tumour heterogeneity. Due to this, recent studies have emerged that have further classified TNBC into four primary subtypes, with each having distinct transcriptional programs and differing responses to chemotherapy ^7, 42^. To address this and explore the role of our 20 gene panel in each subtype we performed pseudobulk RNA-seq analysis, on a dataset containing 6 TNBC patients^43^. We successfully classified each patient into TNBC subtypes; basal-like (BL1 and BL2), luminal androgen receptor (LAR) and mesenchymal (M) using TNBCtype^6^ (Supplementary Fig 4A-B). To broaden our model’s applicable strength, we applied our prediction model to the pseudobulk data, resulting in the prediction of three patients as having a potential for developing RD (Supplementary Fig. 4C-D). Using the R package “UCell”^44^ we measured the average expression of our 20 genes across each subtype and prediction and found that our signature was higher in patients with BL1 and LAR subtypes and patients predicted to have RD (Supplementary Fig. 4E-F). These results suggest that higher expression of a distinct set of genes, originating from specific cellular subpopulations, potentially drives chemoresistance in certain TNBC patients. Overall, our findings revealed a minimalistic gene signature of 20 genes that can predict chemotherapy response in treatment naïve TNBC patients with high accuracy and hold strong potential for prognosis in these patients.

### A distinct epigenetic landscape defines chemoresistance status

Epigenomic dysregulation is known to play a critical role in disease progression in multiple cancers, including TNBC. The acetylation of Lysine 27 at Histone H3 (H3K27ac) is a mark of active proximal and distal regulatory elements including enhancers and known to govern the gene expression programmes associated with cell identity. Therefore, we analysed ChIP-seq) data for H3K27ac for eight primary TNBC patients as well as corresponding transcriptome (RNA-seq) datasets (ENA: accession number PRJEB33558). First, by applying our therapy resistance prediction model to the RNA-seq data from each patient, we were able to classify each as having a potential for developing RD while normal human mammary epithelial cells (HMEC) as pCR (Fig. 4A). We next classified each patient sample into four TNBC subtypes, basal-like (BL1 and BL2), luminal androgen receptor (LAR) and mesenchymal (M) using TNBCtype^6^. Using a similar approach, we also classified TNBC cancer cell lines into TNBC subtypes (Fig. 4A). Furthermore, given the tumour-cell exclusive origin of our signature, we also attempted to classify these cells lines as pCR and RD and were successful (Fig. 4A).

**Figure 4.**
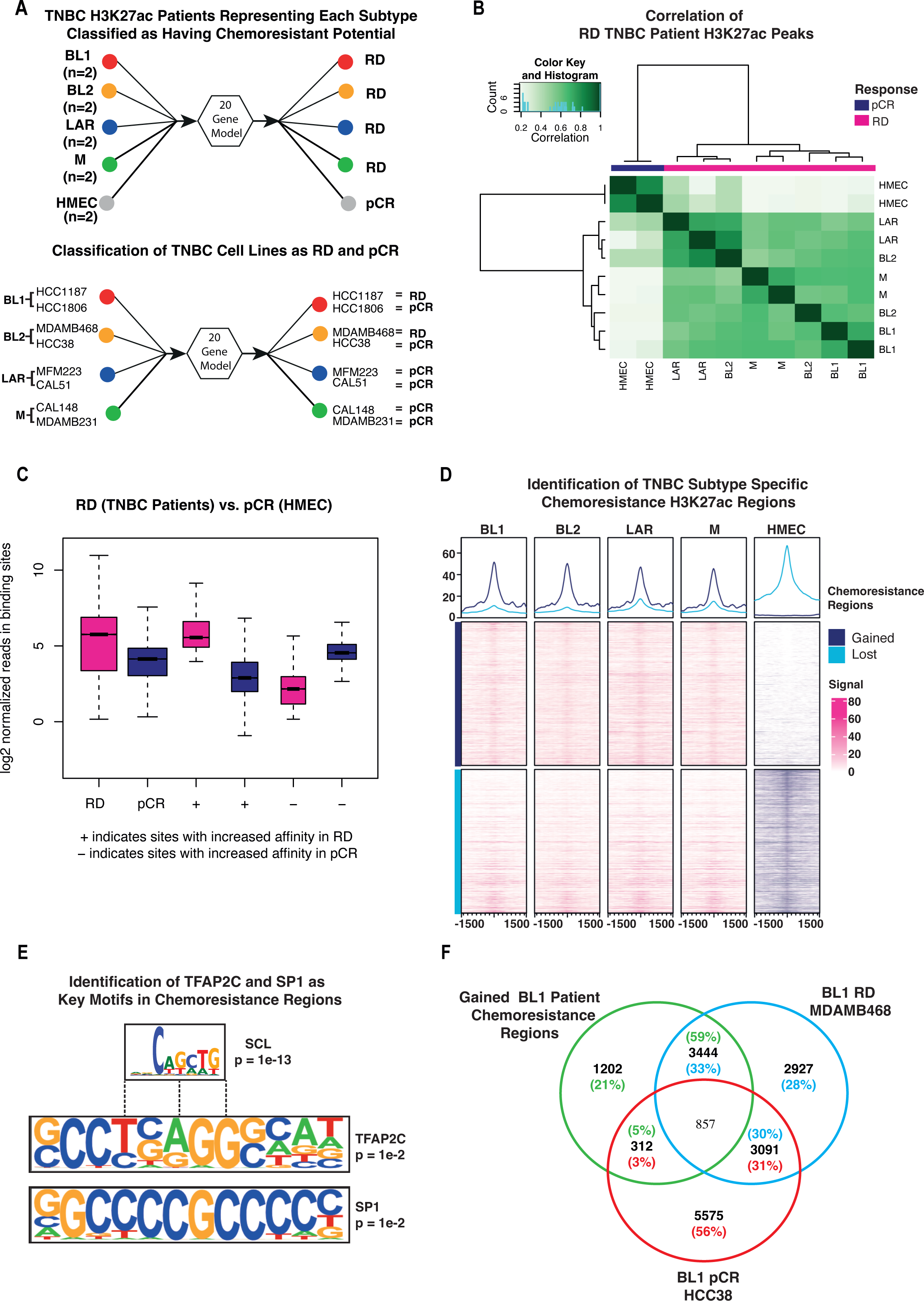
A distinct epigenetic landscape defines chemoresistance status. **A)** Prediction results of 8 TNBC patients and cell lines representing each TNBC subtype. **B)** Correlation of H3K27ac peaks between each TNBC patients (RD) and HMEC (pCR) samples **C)** Identification of RD-specific H3K27ac peaks compared to pCR **D)** Using genomic locations for chemoresistant genes identified in pre-treatment scRNA-seq data we identified regions gained in each TNBC subtype (RD) which are lost in HMEC (pCR) **E)** Identification of TFAP2C and SP1 among key motifs at chemoresistance regions **F)** Comparison of gained chemoresistant gene regions with BL1 RD cell line MDAMB468 and BL1 pCR cell line HCC38.

We next investigated how the distribution of H3K27ac changes across these subtypes. Towards this, we compared the overlap of all H3K27ac peaks across each TNBC subtype and identified regions with an increased enrichment in RD compared to pCR (Fig. 4B-C). Next, to characterise the chemoresistance genes using subtype-specific H3K27ac signal, we compared the loss and gain of H3K27ac marks within chemoresistant gene regions across each subtype and pCR samples. Differential enrichment of peaks between pCR and RD was called using the R package “Diffbind”^45^, with the criteria of a minimum of 50% overlap of peaks. This analysis further showed that there was a significant gain of H3K27ac enrichment at chemoresistance genes across each TNBC sample (RD) and a significant loss in pCR (Fig. 4D). However, this analysis was not able to fully discriminate regions that gained sites (with positive fold change) and lost sites (negative fold change) when comparing pCR and RD. As we had used these criteria to call these peaks (differential enrichment), it is possible that some regions still had sufficient enrichment of H3K27ac that shows up in the heat map even though it is reduced in comparison to the other condition.

To gain insights into the regulatory machinery, we next sought to identify binding sites of specific transcription factors at chemoresistance genes with acquired H3K27ac marks in RD patients. Motif analysis revealed again a strong enrichment for TFAP2C and SP1 motifs among others (Supplementary Table 3), with strongest regulon activity in chemoresistant patients (Fig. 2B-E), clearly implying them as potent drivers of the chemoresistance state (Fig. 4E). Furthermore, chemoresistance genes that gained H3K27ac in BL1 RD patient data showed a stronger overlap with similar genes in the RD cell line compared to the pCR (Fig. 4F), showing a conserved nature of contribution of these genes and their upstream regulation in chemoresistance across systems.

### Unique super-enhancers are associated with TNBC-subtype-specific transcriptional programs underlying chemoresistance

Whilst we were able to show that many chemoresistant genes had strong H3K27ac signal in TNBC patients predicted to have RD, this was not sufficiently discriminatory (Fig. 4D). We, therefore, focused on the analysis of super-enhancers (SEs) which have increasingly been implicated in disease initiation and progression in various contexts including cancer^15^ ^46, 47^. This is particularly interesting as no studies have yet investigated their contribution to TNBC chemoresistance. We, therefore, subjected our genome wide H3K27ac profiles for TNBC patients to the identification of SE elements. SEs were mapped and quantified by Rank Ordering of Super-Enhancers (ROSE) software. In summary, ROSE analysis was performed with default parameters of 12.5 kb stitching distance, and TSS exclusion size set to 0, with the genome set to hg38 ^48^. SE-associated genes were identified as the “nearest gene” output from ROSE. Samples were merged based on their subtyping, to identify common subtype-specific SE regions (Fig. 5A), resulting in an average of 1279 SEs identified per tumour sample (Fig. 5B). The genome wide distribution of H3K27ac SE peaks showed its distribution mostly at intron (59%) and intergenic (33%) locations (Supplementary Fig. 5A).

**Figure 5.**
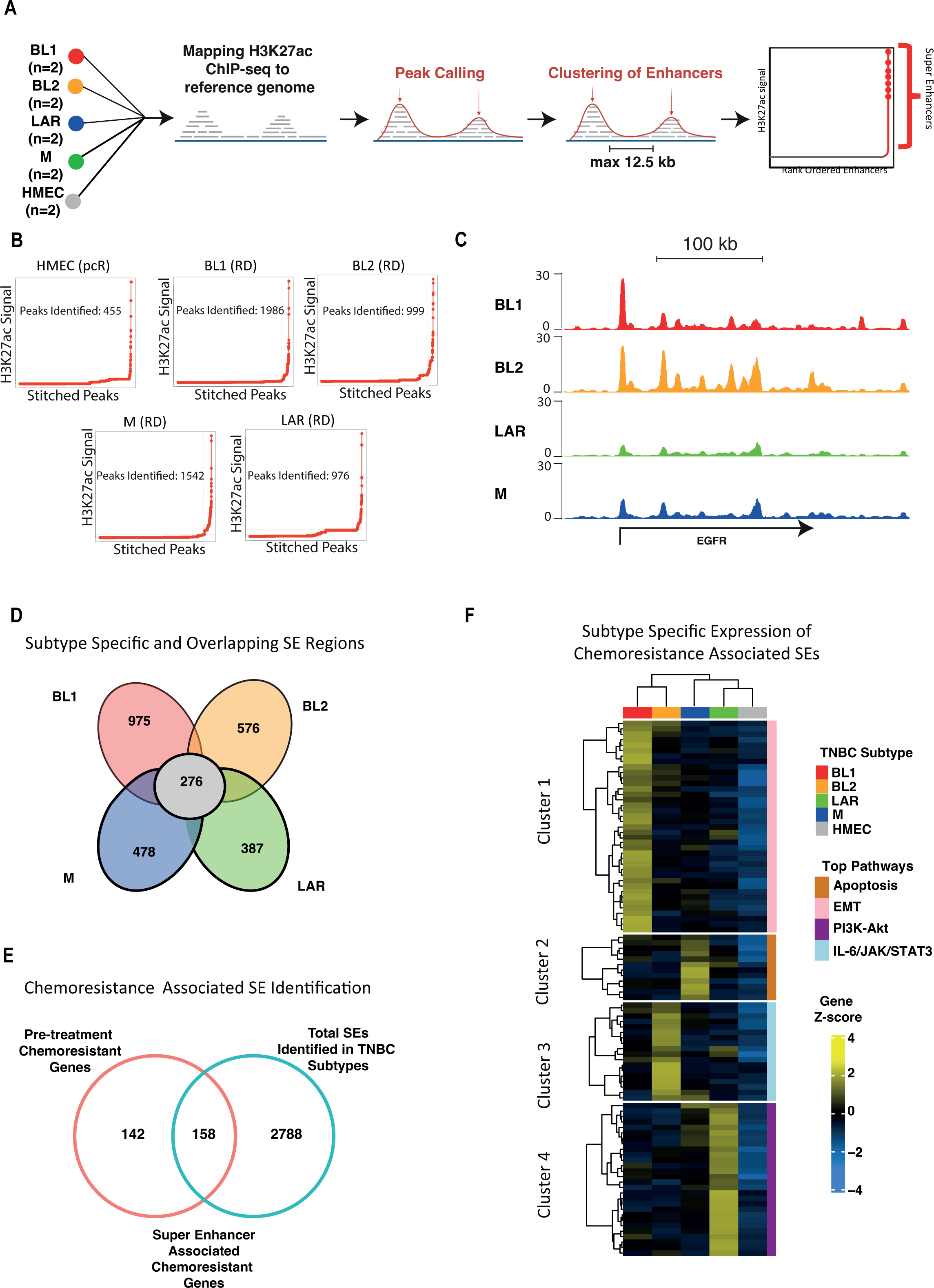
Epigenomic profiling reveals super enhancers in TNBC subtypes which drive the expression of chemoresistance markers. **A)** Schematic showing key steps for Super Enhancer identification **B)** ROSE output of total super enhancers identified in each TNBC subtype **C)** Genome browser track of EGFR, a super enhancer identified in BL1 and BL2 samples **D)** Identification of unique and overlapping super enhancer regions across each TNBC subtype **E)** Identification of SEs overlapping with chemoresistance genes identified in our reproducibility analysis **F)** Heatmap of SE associated genes unique expression between TNBC subtypes. Unsupervised hierarchical clustering demonstrated clustering of genes into 4 clusters, demonstrating unique expression of SEs within each TNBC subtype.

In the first instance TNBC SE regions were compared with HMEC pCR samples to identify TNBC-specific SE regions (Supplementary Fig. 5B). Further analysis of these data identified 2692 unique and 276 overlapping SEs between each TNBC subtype (Fig. 5C-D). We hypothesised that these subtype-specific SEs govern the expression of a selected set of our marker resistance genes to drive TNBC chemoresistance. Interestingly, of our 300 marker genes, 158 were in close proximity to the discovered subtype-specific SEs (Fig. 5E). Next, we calculated the correlation of expression of SE-associated genes across all patients and performed unsupervised hierarchical clustering to identify SE-associated genes that show subtype-specific expression (Fig. 5F). This analysis identified four prominent clusters with unique characteristics of each subtype, and which showed no expression in HMEC cells. Interestingly, further Gene Ontology analysis showed enrichment of specific pathways in each of these subtype-specific clusters. For example, cluster 1, consisting of BL1 specific SEs, was enriched with EMT related signatures; cluster 2, consisting of M specific SEs, was associated with Apoptosis related signatures; cluster 3, consisting of BL2 specific SEs, was enriched for IL-6/JAK/STAT3 signalling signatures while cluster 4, consisting of LAR specific SEs, showed PI3K-Akt related signatures (Fig. 5F, Supplementary Fig. 5C). Of note, EGFR and RAI1, two markers we identified to have a high discriminatory effect in chemoresistant patients, were located in close proximity to a distinct set of discovered SEs in BL1 and BL2 subtypes. Furthermore, our analysis showed that the subtypes LAR and BL1 exhibit the highest number of chemoresistant SEs (Fig. 5F). These findings are in line with previous research, where LAR followed by the BL1 subtype showed the worst response to chemotherapy ^49^. To confirm that these SE regions were communicating with the predicted target chemoresistance genes we processed existing Hi-C datasets from TNBC patients^50^. By searching the SE regions output by ROSE, we indeed confirmed that many SEs of interest are looping in close physical proximity to their predicted target genes, including EGFR and RAI1 (Supplementary Fig. 5D). Altogether, these observations suggest that the super-enhancer landscape plays a key role in the evolution of chemoresistance by governing the expression of key driver genes/pathways in a TNBC subtype-specific manner.

### Distinct transcription factor core regulatory circuitries operate at TNBC subtype-specific super-enhancers associated with chemoresistance

Super Enhancers recruit a high density of cell type-specific master TFs to drive cell-state-specific gene expression profiles^51^. Furthermore, the expression of TFs that bind SEs are often regulated by the activity of SEs in a forward feedback loop and is well-established in many malignant cell types ^33–34^. To reveal critical master TF interactions responsible for driving the TNBC subtype-specific transcriptional program associated with chemoresistance, we modelled transcriptional regulatory networks mediated by SEs utilising the python package “CRCmapper”. It scans TF motifs inside chemoresistance SE regions, and then identifies both TFs binding within SE regions and outward binding of SE-associated genes in a complete regulatory circuitry (Core Regulatory Circuitry (CRC) cliques). CRC cliques are then scored based on TFs which exhibit a high frequency of occurrence across each CRC clique, and the top CRC for each TNBC subtype is designated (Fig. 6A).

**Figure 6.**
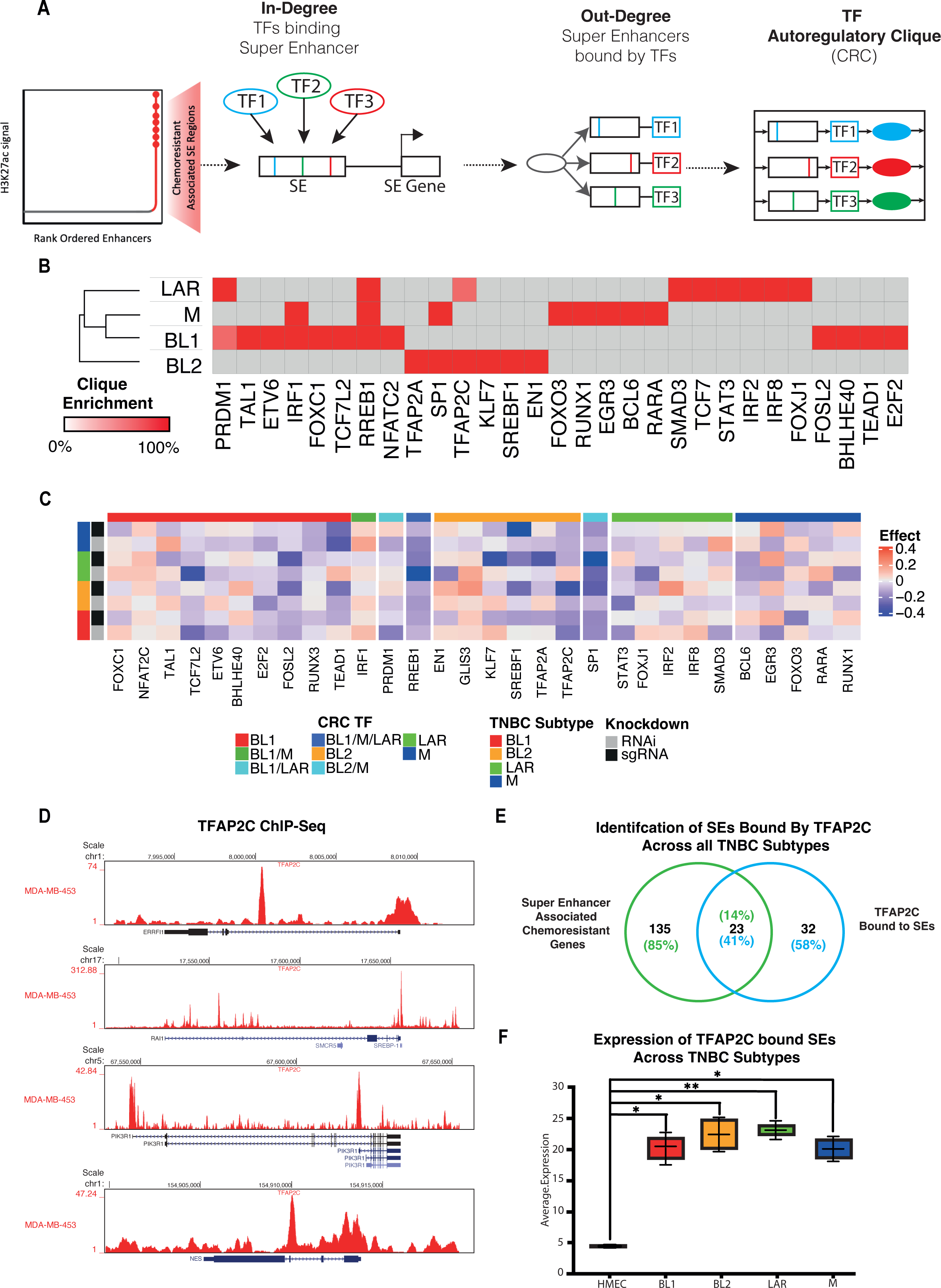
SE - TF connectivity analysis defines core regulatory circuitry underlying TNBC chemoresistance. **A)** Schematic of SE-based CRC analysis. For every TF associated with a chemoresistant SE, in degrees values are calculated by motif identification; Out degrees values are calculated for each TF associated with a chemoresistant SE by determining all other bound SEs at each TF gene loci. Node connections between TFs are used to identify auto-regulatory cliques that in turn regulate the chemoresistant SE network **B)** A heatmap of clique enrichment scores for the union of all TFs associated with top SEs across all TNBC samples. Gray boxes are used when a TF is not associated with a TNBC patient sample. TFs and samples are clustered by Euclidean distance. **C)** Subtype-specific genetic dependencies of each CRC TF from Broad DepMap whole-genome RNAi and CRISPR screen. Heatmaps show significant subtype-specific genetic dependencies using a modified T-test corrected for multiple hypothesis testing (T-value, FDR <0.1). **D)** UCSC Visualization of TFAP2C occupancy using the ChIP-seq data at SEs predicted to be regulated by TFAP2C **E)** Identification of TFAP2C bound SEs across TNBC subtypes **F)** Expression of TFAP2C bound SEs across each TNBC subtype and HMEC.

Following “CRCmapper” analysis on each sample, we calculated a clique enrichment score (the percentage of each CRC in which a TF is a constituent member) (Fig. 6A). Following the scoring of each CRC, we clustered samples based on their clique enrichment scores and revealed intrinsic CRC differences in top TFs across TNBC subtypes and additionally highlighted overlapping TFs common between all subtypes (Fig. 6B, Supplementary Fig. 6A-B). The TFs identified in each TNBC CRC clique include known lineage-defining TFs such as EN1 ^56^. Other TFs, with a high clique score, include EGR3, STAT3, ETVS, IRF1, IRF2, and IRF8. Additionally, we performed CRC analysis using HMEC SE data to identify the top CRC in normal (pCR) (Supplementary Fig. 6C). Of note, FOXC1 was also discovered among other TFs to be a CRC TF in BL1 in line with the recent findings of it being a SE master regulator of invasion, metastasis and chemoresistance in TNBC ^57, 58^.

We next sought to explore the role of identified TNBC-subtype CRC TFs in exhibiting strong genetic dependency across multiple TNBC subtypes. Through this analysis we sought to identify possible candidate TFs that could be implicated, across subtypes, in driving chemoresistance and hold the potential for novel therapeutic intervention. By analysing viability data, (following RNAi and CRISPR knockdown) from the Broad DepMap project, we extracted data for each CRC TF from cell lines representing each TNBC subtype. We performed a regression analysis to associate the correlation between cell line viability with each subtype following the loss of function assays. This analysis revealed CRC TFs which negatively and positively affected the viability of each TNBC cell line and identified key genetic dependencies specific to each subtype and across all (Fig. 6C). Of note, TFAP2C and SP1 were discovered to be essential for viability across all TNBC subtypes. Other TFs, such as STAT3, show genetic dependency across all subtypes, however in some subtypes, they are stronger when compared to others possibly due to their engagement in other networks (Fig. 6C). Furthermore, RREB1 has been shown to be a critical integrator of TGFβ and Ras signalling pathways during both developmental and cancer EMT programs ^59^.

Given our previous observations for a strong enrichment of TFAP2C motifs at the super enhancers associated with chemoresistance genes (Supplementary Table 3), its very high regulon activity and expression in chemoresistant patients (Fig. 2B-E) and now a strong genetic dependency in TNBC cells, we were very tempted to further investigate TFAP2C function in driving TNBC chemoresistance. Using the Super-Enhancer Archive^61^ we first identified SEs and their corresponding TFs and overlapped with our CRC and ROSE analysis outcomes, which further resulted in a putative list of top CRC TFs predicted to be bound by TFAP2C. To identify direct targets of TFAP2C, we processed TFAP2C ChIP-seq from the TNBC cell line MDA-MB-453^60^. A visualization of TFAP2C binding at predicted SE regions indeed confirmed its strong enrichment at these locations (Fig. 6D). Of note, the SE RAI1, a top chemoresistance signature gene in our prediction model, showed a significant occupancy by TFAPC. We next overlapped all SEs bound by TFAP2C across TNBC patients (n=55) with all SE regions associated with chemoresistance genes, resulting in a total of 23 chemoresistance SEs occupied by TFAP2C (Fig. 6E). Notably, these loci included our chemoresistance signature genes as well as other potentially interesting candidates (examples shown in Fig. 6D). Interestingly, gene expression analysis of chemoresistance genes associated with TFAP2C bound superenhancers showed that they were expressed at significantly higher levels in all TNBC subtypes as compared to the healthy control cells (HMEC) (Fig. 6F). Altogether these observations highlight that distinct transcription factor CRCs operate at TNBC subtype-specific super-enhancers associated with chemoresistance genes and TFAP2C holds potential as one of the key TFs of this process across all TNBC subtypes.

### TNBC-type specific CRC TFs are essential for the viability of TNBC cells, and their loss enhances sensitivity to chemotherapy

We next sought to experimentally investigate whether the predicted CRC TFs actively control the expression of chemoresistance genes and consequently chemotherapy response. Our results revealed that TFAP2C is potentially a master regulator across all TNBC subtypes in driving chemoresistance genes by targeting their SEs. Furhermore, equally interesting was SP1 that similarly also showed a strong enrichment at SEs of chemoresistance genes and high regulon activity and expression in chemoresistance cells. We therefore performed depletion of TFAP2C and SP1 in four TNBC cell lines representing each TNBC subtype and measured expression of target SE associated genes using RT-qPCR assays (Fig. 7A & Supplementary Fig. 6D). Interestingly, in all types, knockdown of TFAP2C and SP1 led to a significant decrease in the expression of genes associated with their target chemoresistance SEs (Fig. 7B).

**Figure 7.**
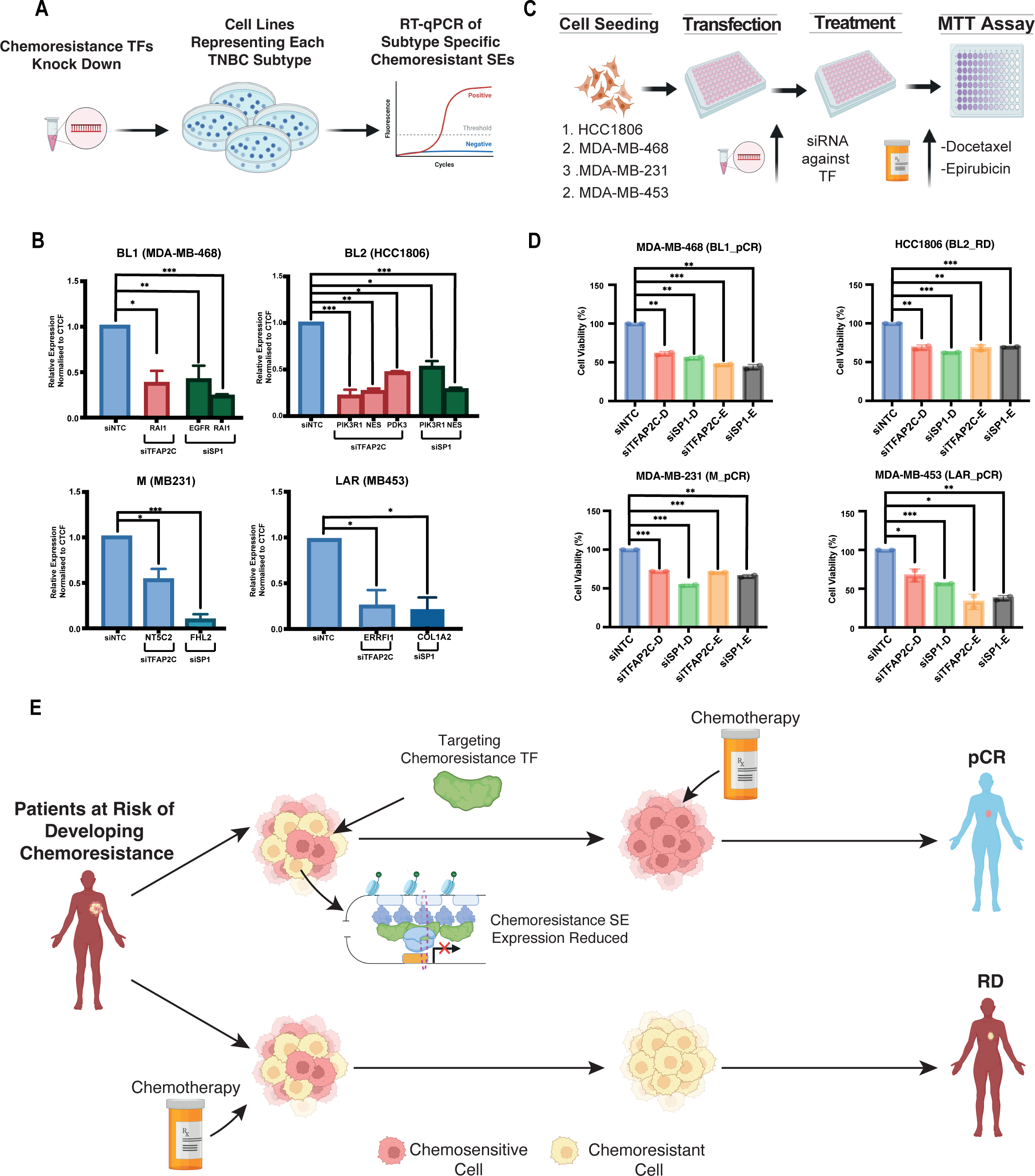
TNBC-type specific CRC TFs are essential for TNBC cell survival, and their depletion improves chemotherapy response. **A)** Schematic of depletion of selected TFs in cell culture followed by RT-qPCR experiments. Workflow for the depletion of CRC TF in selected cancer cell lines. **C)** RT-qPCR of subtype specific chemoresistant SEs following KD of TFAP2C and SP1 (Student’s *t*-test). **D)** Reduced cell viability following chemotherapy treatment in TNBC subtype specific cell lines when combined with depletion of the TFs (Student’s *t*-test). **E)** Targeting chemoresistance SE associated TFs has the potential to eradicate subpopulations associated with chemoresistance and improve chemotherapy efficacy.

Since we had found that many SEs were bound by TFAP2C and SP1, and have a strong genetic dependency across each TNBC subtype, we next investigated whether such loss in the expression of chemoresistance genes following their depletion would improve the response to chemotherapy. Therefore, we performed siRNA knockdowns of each candidate TF in four cell lines representing each TNBC subtype, accompanied by treatment with two chemotherapy agents (Docetaxel and Epirubicin) and measured cell viability using an MTT assays (Fig. 7C). Since these TFs may have other physiological roles ^35, 56, 62–65^ in TNBC beyond the context of chemotherapy, we did not characterise the effect of depleting these TFs alone, without the drugs. Interestingly, the depletion of TFAP2C and SP1 led to a significant decrease in cell viability following chemotherapy treatment in all TNBC subtype cell lines compared to the control cells (Fig. 7D). In particular, TFAP2C knockdown showed a significant reduction in cell viability across all subtypes, again highlighting that TFAP2C is a master, versatile regulator of chemoresistance across all TNBC subtypes. Overall, our results show that a distinct set of TFs potentially drive the epigenomic landscape and govern the gene expression programme that define chemoresistance-associated cell subpopulations. Furthermore, direct targeting of these chemoresistance-associated TFs holds the potential to improve patient outcomes across all TNBC subtypes (Fig. 7E).

## DISCUSSION

Neoadjuvant chemotherapy (NAC) is used frequently in the treatment of TNBC patients due to the lack of targeted therapeutics and its ability to reduce tumour size, improve surgical outcomes and increase survival in responders. However, due to the intratumoral heterogeneity (ITH) associated with TNBC, patients have differing responses to NAC ^66^. Achieving pCR is associated with significantly improved survival outcomes in TNBC patients ^67^. Identifying those patients who will have RD following NAC will enable physicians to determine the best therapeutic option at the beginning of treatment, rather than waiting for NAC treatment results, to increase the chances of achieving pCR. Numerous efforts have been put into developing predictive signatures in TNBC, but currently, there is no clinically recommended predictive biomarker panel for NAC response ^8, 68, 69^. However, these studies have focused on bulk RNA based techniques, in small patient cohorts to identify markers to predict therapy response and do not account for the ITH associated with TNBC.

Here, by profiling chemoresponsive and chemoresistant patients at the single-cell level to identify markers associated with chemotherapy, we have developed a predictive model which has high accuracy in defining chemotherapy response in TNBC patients. Our 20 gene model, through the identification of markers in scRNA-seq and validation in over 300 patients, holds a high potential for aiding in the clinical management of TNBC patients by enabling the assessment of NAC response upfront. Additionally, we have demonstrated that it outperforms all existing signatures for predicting chemotherapy response in TNBC. Furthermore, higher expression of our models’ genes are also associated with reduced survival and could accurately predict the chemoresistance potential in TNBC patients from the METABRIC cohort. It is strongly associated with EGFR signalling, which has been shown to play a critical role in TNBC chemoresistance ^26–28^.

The predictive strength of our model’s combination of genes was further demonstrated by predicting chemotherapy response in the eight untreated TNBC patients with H3K27ac data. Our model was able to classify the patients as having potential for RD accurately. This provided the unique opportunity and the rationale to map and quantify enhancers genome-wide to shed light on the previously uncharacterised SE landscape underlying chemoresistance in a subtype dependant manner for the first time. By overlapping with chemoresistance genes identified to have a reproducible expression in bulk RNA data, we could identify a subset of chemoresistance genes in close proximity to SEs. Interestingly, BL1 and LAR subtypes had the highest proportion of SEs. LAR followed by BL1 are the top two TNBC subtypes associated with increased chemoresistance and poorer outcomes ^49^. The SE-associated genes were significantly involved in EMT and PI3K-Akt signalling processes in BL1 and LAR subtypes. Both have previously been implicated in chemoresistance in multiple cancer types, including breast cancer and specifically TNBC ^70, 71^.

Additionally, whilst the dysregulation of gene expression in TNBC has been previously characterised, it has not been adequately explained by typical transcriptomic analysis alone. Instead, such extensive changes are attributed to the widespread transcriptional rewiring occurring in breast cancer cells, including the utilisation of core transcription factors, as well as the activation of many gene-regulatory elements, including enhancers and super enhancers.^13, 72^ Defining epigenomic characteristics are instrumental to dissecting gene regulatory programs which underlie cancer disease progression. Here, for the first time, we have profiled and characterised the TF regulatory network, using subtype-specific SE profiles, underlying TNBC chemoresistance. This systematic identification of active TNBC subtype regulatory elements has led to several enabling observations. By constructing the TNBC TF regulatory network using subtype-specific SE profiles, we can identify the critical TF nodes that enforce the TNBC subtype epigenome underlying chemoresistance. Of note, TFAP2C, TFAF2A, and SP1 were shown to have higher expression in chemoresistance pre- and post-single-cell data, highlighting their implication in driving chemoresistance in TNBC. Additionally, by profiling TFAP2C ChIP-seq in a TNBC line line we found that many chemoresistance associated SEs, including RAI1, were bound by TFAP2C, establishing its direct function in driving TNBC chemoresistance. Furthermore, we depleted key chemoresistance TFs predicted to function at subtype-specific chemoresistance SEs of chemoresistance genes and measured their expression using RT-qPCRs (Fig. 7B). These results show a clear, significant reduction in the expression of target chemoresistance genes, validating our proposal for the role of these TFs in regulating their expression. These results are also in line with our observations that the depletion of these TFs can significantly overcome chemoresistance (Fig. 7D). Altogether, these observations conclude that chemoresistance is governed by a distinct set of genes that are controlled by CRC TF networks through a subtype-specific set of SEs.

Additionally, they were shown to have a high genetic dependency in each TNBC subtype cell line. Of note, suggesting that inhibition may provide an approach to overcome chemotherapy resistance in all TNBC subtype tumours. In recent years, targeting TFs using small molecules which bind to specific nuclear hormone receptors has proven to be successful in many cancers ^73^, in particular the SP1 inhibitor Mithramycin A has been shown to inhibit and suppress cell survival in *in vitro* models of basal TNBC ^74^. Along these lines, targeting TFAP2C may dramatically improve chemotherapy efficacy in patients with a high risk of chemoresistance.

In BL1, FOXC1 was highlighted as one of the top TFs driving chemoresistance SEs. In the single-cell data, FOXC1 was shown to have higher expression in chemoresistant cells pre- and post-chemotherapy. FOXC1 has recently been shown to be a master TF, encoded by SEs in TNBC ^57^. Additionally, TFAP2C has never been shown to drive SE expression in TNBC, nor has it previously been implicated in TNBC chemoresistance. Its key role in potentially driving TNBC chemoresistance is further highlighted by our SCENIC analysis. We identified TFAP2C, along with several other CRC TFs, as key regulons in defining the chemoresistance subpopulations in the scRNA-seq data. While it was identified as a core TF in BL2, we have demonstrated it has a high genetic dependency and potential regulator of chemoresistance SEs in all TNBC subtypes. Additionally, it has been implicated in chemoresistance in several cancers ^31, 75^ and, notably, Docetaxel resistance in lung adenocarcinoma ^76^. The TFs TFAP2C and SP1 were identified throughout our study from the single-cell to CRC analysis as potentially having a significant role in driving chemotherapy resistance-associated gene expression programme. We have successfully shown that direct targeting of these TFs has the potential to increase the efficacy of chemotherapy agents across each TNBC subtype. Whilst there are no clear TFs that act in a unique subtype dependant manner, we have shown that the TF TFAP2C is a master regulator of subtype-specific chemoresistance SEs across all TNBC subtypes resulting in potential for the development of novel therapeutics that can aid in improving the efficacy of NAC.

Our results have clearly highlighted how a better understanding of gene regulatory circuitry allows identifying novel therapeutic avenues. This study creates the rationale for further functional studies to determine their mechanistic roles in chemoresistance and potentially lead to the development of novel targeted therapeutics. Additionally, as the model was developed based on a combination of NAC, it may be possible to extend its application range to develop drug-specific or secondary therapeutic prediction models and further stratify TNBC patients. One potential limitation of our study is the low number of patient samples for SE identification. However, the genes identified in close proximity to SEs were shown to have higher expression in RD TNBC patients across multiple studies and TNBC subtype-specific cell lines, validating their role in TNBC chemoresistance.

In summary, we reveal cell subpopulations associated with TNBC chemoresistance and the signature genes defining these populations of which a subset acts as a best-in-class gene signature for an accurate prediction of chemotherapy response. Notably, we show that these chemoresistance genes are controlled by a specific set of transcription factor networks and super-enhancers in a TNBC-subtype specific manner. Importantly, we demonstrate that targeting these TFs holds the potential to overcome chemoresistance and ultimately improve patient survival.

## Acknowledgements

We would also like to thank the members of the Tiwari lab for their cooperation and critical feedback throughout this study. We would especially like to thank Mohammed Inayatullah and Rasmus Siersbæk for their critical comments on the study. The support from the Core Facilities of the Queen’s University Belfast and University of Southern Denmark is gratefully acknowledged. This study was supported by the Deutsche Forschungsgemeinschaft TI 799/1-3, Innovation to Commercialisation of University Research (ICURe) and Department for the Economy (DfE).

## AUTHOR CONTRIBUTIONS

R. L. and V.K.T. designed the study, analyzed data, and wrote the manuscript. Z.Z. and A. M. performed experiments.

## DATA AND CODE AVAILABILITY

The merged microarray datasets and all scripts used in this study are located at: <u>Github.com</u> Accession numbers for all publicly available datasets used are in: Supplementary Table 1 Sheet 1

## SUPPLEMENTARY TABLES

**Supplementary table 1:**

Sheet 1: Accession numbers of public datasets used in this study. Sheet 2: Cell type markers used for annotating the scRNA-seq data. Sheet 3: The list of 300 Chemoresistant Genes.

Sheet 4: Cell Line Classification.

Sheet 5: Sequences of siRNAs used in this study. Sheet 6: Sequences of primers used in this study.

**Supplementary table 2:**

Regulons identified in the SCENIC analysis.

**Supplementary table 3:**

Motifs enriched at H3K27ac sites.

## METHODS

### Identification of chemoresistant cell types using single-cell RNA-sequencing analysis

To identify cell types and their markers associated with TNBC chemoresistance, scRNA-seq analysis was performed on the data set was obtained from Kim et al ^20^, consisting of matched pre and post-chemotherapy (anthracycline and a taxane) samples from four responsive and four resistant patients with a total of 6,862 cells. To integrate the single cell data from each patients’ samples, functions FindIntegrationAnchors and IntegrateData from Seurat v3 were implemented. The integrated data was then scaled and downstream analysis, including normalisation, variable feature selection, dimensionality reduction and UMAP clustering, was performed. Cluster annotation was performed using the python program “SCSA” to identify associated cell types and cancer-related processes. All significantly expressed markers for each treatment time point and therapy response were polled to identify uniquely expressed markers in pre-chemoresistant patients that could potentially have a crucial role in driving TNBC chemoresistance.

### Reproducible Signature Marker Identification

For the reproducibility of the gene set identified using scRNA-seq analysis, we used five independent bulk RNA-seq datasets, GSE20271, GSE25055, GSE25065, GSE20194, GSE163882, of 397 TNBC patients where their chemotherapy response was available (RD, PCR). Patient samples were excluded if the therapeutic outcome (residual disease or pathologic complete response) was unknown and not classified as TNBC. The raw data was normalised, batch corrected, and log-transformed using the R package “affy” and “TDM and the python package “pyComBat”. In total, 397 patient’s data were selected for reproducibility analysis. The 300 genes identified in the scRNA-seq dataset were extracted from the normalised bulk RNA-seq count files. A custom R script was used to compare each gene’s expression in patients with residual disease and pathologic complete response, Wilcoxon Rank Sum and Kruskal-Wallis tests were used to calculate significance.

### Pseudobulk Analysis

All six patient data files were downloaded from: and GSE118390 and the chemoresistant scRNA-seq were analysed using the same parameters in the R package “Seurat”. First, cells with feature counts of greater than 2500 or less than 200 were removed, including mitochondrial reads of greater than 5%. Following the removal of cells, downstream analysis, including normalisation, variable feature selection, dimensionality reduction and UMAP clustering, was performed. Signature scoring was performed by the R package UCell^44^ using default parameters. Following downstream analysis, pseudobulk analysis was performed using the R package SingleCellExperiment and the function “AggregateExpression”.

### Implementation of GENIE3 and SCENIC

Single-Cell regulatory Network Inference and Clustering (SCENIC) analysis was performed to reveal the core TFs in chemoresistant and chemosensitive clusters^29^. We performed the SCENIC analysis using the latest version of pySCENIC. The gene-motif rankings (500 bp upstream or 100 bp downstream of the transcription start site) were used to determine the search space around the TSS. The motif database was used for RcisTarget and GENIE3 algorithms to infer the core TFs. Wilcoxon Rank Sum and Kruskal-Wallis tests were used to calculate significance.

### Identification of Significant Gene Set and Construction of the Prognostic Prediction Model Based on Residual Disease Vs Pathologic Complete Response

Raw microarray expression (CEL) files of all 310 TNBC patients were downloaded from Gene Expression Omnibus, GSE20271, GSE25055, GSE25065, GSE20194. Gene expression profiles were quantile normalised and log2 transformalised using “BART”, followed by batch correction using “ComBat” from the R package “sva”. To identify the most significant gene set, GSE20271 and GSE25055 datasets with 177 TNBC patients (57 pathologic complete response, 120 residual disease) were used to build the model. To verify the strength of the geneset, GSE25065 and GSE20194 datasets with 130 TNBC patients (46 pathologic complete response, 84 residual disease) were used as the external validation cohort. To identify the significant gene set and develop a predictive model to discriminate pCR and RD groups, we first used Lasso and Elastic-Net Regularized Generalized Linear Models using the R package “glmnet” on the 300 markers to identify the best combination with the greatest predictive power. Then, we used the 10-fold cross-validation method to evaluate the discrimination ability, between pCR and RD, to obtain a relatively unbiased estimate. After the LASSO regression analysis, a predictive model based on 20 was used to fit a generalised linear model. The predictive capability was measured by the receiver operating characteristic curve (ROC curve) area under the curve (AUC) using the R package “pROC”. Results were evaluated using the area under the ROC curve. The optimal model was selected by maximising AUC. The model was tested on data with known and unknown chemotherapy response using the function predict.glm with the <u>ideal</u> lamda as the s variable.

### TNBC subtyping

The TNBCtype web-based tool (http://cbc.mc.vanderbilt.edu/tnbc/) was used to classify each TNBC patient sample. Subtyping was performed on RNA expression data, normalised within TNBC patients as recommended by the tool, from each patient.

### H3K27ac ChIP-seq Analysis

Eight primary TNBC patients as well as corresponding transcriptome (RNA-seq) datasets were downloaded from ENA: accession number PRJEB33558. Reads were aligned to the human genome (GRCh38) using Bowtie2. H3K27ac ChIP peaks were identified by the MACS version 2 software package with paired input samples with the callpeak function using default settings, genome set to ‘hs’, and peak calling set to—broad. Differential enrichment of peaks between pCR and RD was called using the package diffbind with the criteria of a minimum of 50% overlap of peaks.

### Super Enhancer Identification and Analysis

Samples were merged based on their subtyping using bedtools merge and enhancer and SE elements were mapped and quantified by MACS and ROSE software^48^. ROSE analysis was performed with default parameters of 12.5 kb stitching distance, and TSS exclusion size set to 0, consistent with prior studies, we did not exclude TSS elements^77^. Using the output SE bed file from ROSE we identified regions unique to each TNBC subtype using ChIPpeakanno with 50% overlap of SE regions. SE-associated genes were identified by ROSE by assigning the discovered SEs to the nearest genes. Hierarchical clustering on SE-associated uniquely expressed genes was performed using Euclidean distance metric and Ward’s linkage method and plotted using the R package “ComplexHeatmap”. Colour bars for associated pathway data for each subtype were determined using EnrichR.

### Hi-C Analysis in TNBC patients

Samples where obtained from GSE167150 and processed using HiCExplorer^78^. Reads were aligned to the human genome (GRCh38) using Bowtie2. Then the HiCExplorer pipeline was implemented with default parameters.

### TNBC Chemoresistance CRC Reconstruction

We performed the core transcriptional regulatory circuitry analysis using CRC mapper (https://github.com/linlabcode/CRC) as previously described ^79^. Within Super Enhancer regions, the CRC software uses FIMO to find enriched (*q* value < 1e−5) TF motif occurrences. CRC first identified TFs that are active, regulated by a proximal, overlapping, or the closest SE region. The total degree is a measure of how often a given TF participates in a regulatory interaction with other TFs. It is defined as the number of unique TFs participating in a regulatory interaction that affects a given TF plus the number of unique TFs that are regulated by a given TF.

### Genetic Dependency

Gene expression for each CRC TF was extracted from TNBC cancer cell lines from CCLE in DepMap (https://depmap.org/portal/). To identify genetic dependencies of subtype-specific CRC TFs, Achilles gene effect scores and dependency scores were downloaded for each subtype TNBC cell lines screened by RNAi and CRISPR from DepMap. We built linear-regression models of each TFs correlation strength and viability each subtype across all TNBC cell lines tested. T-statistic testing was used to evaluate association strength between subtype correlation strength and viability.

### Cell culture

The TNBC lines HCC1806 and HCC70 were maintained in RPMI 1640 (Gibco, 21875034) medium supplemented with 10% FBS, 1% glucose, 1mM sodium pyruvate (Thermo, 11360070). MDA-MB-468, MDA-MB-453, and MDA-MB-231 cells were maintained in DMEM (Dulbecco’s modified Eagle’s medium) with 10% FBS. Cells were grown as monolayers at 37°C in humidified CO2 (5%) incubator.

### siRNA transfection

The scrambled siRNA control and ON-TARGETplus SMARTpool siRNA targeting human TFAP2C, TFAP2C, SP1, STAT3, TCF7L2, PRDMI, and FOSL2 were purchased from Dharmacon. Transfection was performed using Lipofectamine™ RNAiMAX (Invitrogen, 13778150) according to the manufacturer’s instructions. In brief, cells were seeded at 180k/well for MDA-MB-231, MDA-MB-453, HCC1806 and HCC70. Cells were seeded at 250k/well for MDA-MB-468 cell lines. All cells were seeded the day before the transfection. siRNA at a final concentration of 5 pmol was diluted in 45 μL of Opti-MEM (Gibco, 31985047) and 2.25 μl of Lipofectamine RNAiMAX was diluted in 45 μl of OPTI-MEM. The diluted siRNA and Lipofectamine RNAiMAX were mixed and incubated at room temperature for 10 min. Ninety microliters of transfection mixture were added to each well of 12 well plates. Twenty-four hours later, the transfection cocktail was replaced with complete media for each cell lines.

### RNA isolation and RT-qPCR

Total RNA was isolated from cells in culture using Trizol reagent (Ambion, 15596018) according to the manufacturer’s instructions. RNA concentration and purity were measured using the NanoDrop Spectrophotometer. cDNA was synthesised using Verso cDNA synthesis kit (Thermo, 01280858). RT-qPCR was performed in SybrGreen program: 5 min pre-incubation at 95°C; amplification 45 cycles at 95°C for 10 s, 60°C for 10 s and 72°C for 10 s; melting was performed at 95°C for 5 s, 65°C for 1 min, 97°C on hold; final cooling was performed at 40°C for 30 s. Results were analysed and normalised by the relative quantity (ΔΔCt) method. Wilcoxon Rank Sum and Kruskal-Wallis tests were used to calculate significance.

### MTT assay and drug sensitivity analysis

siRNA transfected cells were cultured for 24 h and treated with Epirubicin and Docetaxel at desired concentration for each cell line. DMSO served as vehicle control. The treated cells were incubated for 48 hours, and a cytotoxicity assay was performed using an MTT assay kit (Roche, 11465007001) according to the manufacturer protocol. Briefly, 10 μl MTT (5 mg/ml) was added to each well and allowed to form formazan crystals for four hours in the incubator. 100 μl of solubilization solution was added to each well and incubated overnight in the incubator in a humidified atmosphere. The next day, complete solubilization of the purple formazan crystals was confirmed and then the absorbance values were determined using a microplate reader (BMG FLUOstar Omega) at 590 nm. The experiments were repeated twice, and data are represented as mean ± SD from three technical replicas. Wilcoxon Rank Sum and Kruskal-Wallis tests were used to calculate significance.

### Statistical Analysis

All the statistical analyses were performed using R (version 4.1.1) and GraphPad Prism 9. Student’s *t*-test, Wilcoxon rank-sum test and Kaplan–Meier were utilised in this study. *p*-values of less than 0.05 were considered statistically significant (* = *p* < 0.05; **= *p* < 0.01; ***=*p* < 0.001).

**Supplementary Figure 1. Merged Pre-Treatment Samples and Gene List Selection**

**A)** UMAP with 8 chemoresponsive and chemosensitive patients labelled. **B)** UMAP with cell type annotations. **C)** Expression of NDUFA6 in merged data. (Wilcoxon rank-sum test, p=5.3e-ll) **D)** Expression of NDUFA6 in unmerged data. (Wilcoxon rank-sum test) **E)** GO terms of all markers identified in pre/post chemoresistant and chemosensitive patients.

**Supplementary Figure 2. Quality Control Figures from microarray batch correction**

**A)** Expression values before and after correction. **B)** PCA plots before and after correction

**Supplementary Figure 3. Development of the 20 gene model**

**A)** Selection of the tuning parameter (λ), based on 10-fold cross-validation, using the LASSO model. Vertical lines represent lamda.min and lamda.1se, the red line represents the cross-validation curve and mean binomial deviance against log-λ. **B)** The coefficients of the 20 genes and 21 probe IDs used to construct the predictive model. **C)** ROC curve showing results in the validation cohort (AUC=0.89) **D)** Average expression of the 20 genes in the TCGA-BRCA cohort (Wilcoxon rank-sum test) **E-F)** Survival plot of the 20 genes in ER+ and HER2+ patients from the METABRIC Cohort (Kaplan–Meier) **G)** ROC curve showing the predictive capability following removal of five genes (AUC=0.678). **H)** ROC curve showing the results of 20 iterations of cross validation (AUC=0.90)

**Supplementary Figure 4. TNBC pseudobulk RNA-seq analysis**

**A)** Clustering of untreated TNBC patients. **B)** Patients labelled based on pseudobulk subtype classification. **C)** Classification of RD or pCR of each patient using our gene panel. **D)** UMAP coloured based on our prediction. **E)** Expression of 20 gene panel across TNBC Subtypes (Wilcoxon rank-sum test) and chemotherapy response prediction (Wilcoxon rank-sum test, p = 6.4e-05).

**Supplementary Figure 5. Identification of Tumour Specific Super Enhancers**

**A)** Genomic Distribution of TNBC Super Enhancers. **B)** Venn Diagrams comparing each TNBC subtypes SE with SE’s identified in HMEC samples **C)** Signalling pathways upregulated for subtype specific SEs **D)** Hi-C plots showing loops between SE regions and predicted genes

**Supplementary Figure 6. CRC Analysis reveals highly connected TFs**

**A)** CRC of all TFs identified for each TNBC patient. **B)** CRC common across all TNBC subtypes **C)** CRC of HMEC Samples **D)** Results of RT-qPCR in M and LAR cell lines (Student’s *t*-test).

